# A microfluidic model to recapitulate pH and oxygen gradients in solid tumors

**DOI:** 10.1101/2025.07.05.663284

**Authors:** Yan-fang Li, Leïla Dos Santos, Mahdi Rezayati Charan, Jamie Auxillos, Albin Sandelin, Stine F. Pedersen, Rodolphe Marie

## Abstract

The tumor microenvironment (TME) plays critical roles in cancer development, aggressiveness, and treatment resistance. The TME comprises cellular (stromal) components as well as gradients of physicochemical properties, including hypoxia and acidosis. Understanding of how hypoxia and acidosis gradients impact cancer phenotypes is lacking, in large part due to challenges in precisely mimicking and controlling such gradients in a manner compatible with the growth of cancer- and stromal cells. Here, we design and validate a microfluidic device enabling orthogonal gradients of oxygen and pH. Both gradients are established by diffusion from a nearby source and sink in the observation area in the absence of flow. This produces linear gradients at steady state. Our device enables a wide range of spatiotemporally resolved analyses, from ‘omics to live cell imaging, interrogating the impact of the physicochemical TME on disease development. The design is easily adaptable, making it valuable for a wide range of questions involving physicochemical gradients.

## Introduction

The tumor microenvironment (TME), comprising both its physicochemical and cellular components, is widely understood to play decisive roles in cancer development. As such, the importance of TME hypoxia and acidosis in shaping disease aggressiveness has been demonstrated in numerous cancers, and the development of gradients of oxygen and proton gradients in growing tumors has been shown ^1–3^. The increased availability and resolution of spatial ‘omics analyses such as transcriptomics, proteomics, and metabolomics have revealed the corresponding, profound heterogeneity of the cellular landscape of tumors ^4,56^. While it is thus clear that both exhibit substantial spatial variation, the causal relationships between the physicochemical and cellular landscapes of tumors remain very poorly understood. This, at least in part, reflects the technical challenges of reconstructing a TME *in vitro* that is representative of the complex *in vivo* tumor conditions. For instance, while the responses of cancer- and stromal cells to both hypoxia ^7,8^ and acidosis ^9,10^ have been studied, such studies have largely subjected the cells to one or a few concentrations of one such parameter. However, this completely fails to recapitulate the complexity of the TME, where hypoxia and acidic pH only partially coincide ^111^, and where the specific concentrations, gradients, and combinations of these and other environmental parameters are unknown yet of potentially profound importance for disease development, patient prognosis, and treatment.

While *in vivo* models can mimic patient tumor TMEs well, they do not lend themselves easily to causality analyses regarding the physicochemical TME composition, both because of their complexity and because the multi-parameter live imaging needed to monitor TME gradients is challenging, if not impossible, in such models. On the other end of the spectrum, 2D cell culture models can be controlled but fail to recapitulate the gradients and complexity of patient tumors. In between these extremes, organoid- and complex 3D models offer many advantages but are not ideal for gradient control or for multiparameter live imaging (for a review, see ^4^). To enable the combination of high-resolution live imaging with precise control of microenvironmental gradients, we and others have developed microfluidic tumor models ^12–18^.

Microfluidics devices allow creating and maintaining predictable concentration gradients using laminar flows. Devices made of optically clear materials enable the integration with optical microscopy for monitoring 2- and 3D cell cultures to interrogate phenotypic features such as motility and morphology based on brightfield- and fluorescence imaging. In addition, rapid developments in the sensitivity of spatial ‘omics techniques now make it technically feasible to exploit microfluidics to develop TME models with highly spatially resolved ‘omics information, precise environmental gradients, and reduced reagent volumes and parallelization ^19^. Microfluidics TME models typically use pH gradients to recapitulate TME acidosis conditions. In such devices, a concentration gradient of hydronium ions can be created either using dilution series or diffusion gradients in the absence of convection. In the first case, a pH gradient is generated by first creating a dilution series using a micromixer device ^20,21,22^ and then merging solutions at different pH in a single channel under laminar flow conditions, thereby creating a step-like pH gradient ^20,21^ that cells seeded in the microchannel will be exposed to. In the second case, a smooth pH gradient can be created between a source and a sink typically consisting of a pair of fluidics channels where low and high pH media is continuously supplied. In such a device, a concentration gradient is established via diffusion in the absence of flow. To suppress flow, cells are embedded in a hydrogel in a plane with a source and/or a sink microchannel ^14^, thereby reproducing matrix conditions closer to the microenvironment in vivo while also suppressing flow. In recent work, we established a microfluidic device and a workflow utilizing microfluidics to create a pH gradient over cells embedded in collagen-I ^12^. The pH gradient generated is stable for at least 24 h, and the device format facilitates live cell microscopy and integration with spatial transcriptomics. A reversible sealing design enables the retrieval of cells embedded in a collagen-I gel, mimicking the extracellular matrix. The cell-containing collagen-I hydrogel can be handled as an artificial tissue and processed through an adapted spatial transcriptomics pipeline. This work allowed us to demonstrate that within a 4 h timeframe, a plethora of genes are differentially regulated as a function of the cells’ position in a pH gradient, mimicking those likely to occur in human tumors. These results highlighted the benefit of integrating a microfluidics gradient generating device with cell culture and spatial transcriptomics to study cancer cells.

However, the TME comprises other physicochemical gradients than pH, including oxygen gradients. Most existing TME-modelling microfluidics devices are made of the gas-permeable polymer PDMS. In such systems, CO_2_ and oxygen concentrations can be fixed through equilibration with the atmosphere generated in the vicinity of the device, either in an incubator or an environmental enclosure of a microscope. However, the cells themselves may participate in this equilibrium through the respiration and make the oxygen concentration effectively difficult to predict and depend on diffusion through the device over large distances (mm to cm) compared to the active area where the cells are. Cellular respiration can itself be utilized to create an oxygen gradient in the chip ^15^ however, in most cases, an oxygen sink is created by flowing an oxygen-depleted gas, usually nitrogen ^16,17,13,18^, medium equilibrated with nitrogen ^23^ or an oxygen scavenger solution such as sodium sulfite ^24,25^ allowing to create either a stable oxygen concentration ^13,17^, a spatial gradient ^13,16,24,25,15^ or spatio-temporal gradient ^13,24^. Oxygen gradients have been combined with other types of concentration gradients (e.g., anti-tumor drugs ^25^); however, the combination of pH and oxygen gradients has, to the best of our knowledge, never been demonstrated.

Here, we developed and validated a microfluidic device that enables the orthogonal superposition of pH and oxygen gradients. As tree-like mixer architecture is only suitable to create multiple gradients that are collinear, we established a pH gradient without flow using diffusion through hydrogel barriers created when the PDMS device is reversibly sealed on top of the cell-containing hydrogel, leading to a linear concentration gradient at steady state. The superimposed orthogonal oxygen gradient was introduced by oxygen and nitrogen flowing in pairs of gas channels placed orthogonally to the fluidic channels supplying the media.

We show that both gradients are linear and in orthogonal directions, effectively creating a matrix of all possible combinations of pH and oxygen tension in the pH range 6.5-7.2 and 1.5-18.2% oxygen. We use this device to investigate the influence of pH and oxygen on the motility of MDA-MB-231 human breast cancer cells. Cell tracking from brightfield time-lapse imaging over 22 h shows a consistent increase in cell median speed at increased pH values. Cell motility is also increased from lower to higher oxygen concentration; however, only in data sets where the cells are generally more motile, emphasizing the need for single-cell measurements to understand cancer cell motility in gradients.

## Materials and Methods

### PDMS device fabrication

The Poly(dimethylsiloxane) (PDMS) microfluidic device consists of two layers (Fig. 1a). Each layer was fabricated by PDMS-casting on a mould as previously reported ^12^. Briefly, the mould was prepared using SU-8 photolithography on a silicon substrate and was coated with 1H,1H,2H,2H-perflourodecyltrichlorosilane. PDMS was prepared using a SYLGARD™ 184 Silicone Elastomer Kit, with monomer and crosslinker mixed at the weight ratio of 10:1. The mixture was degassed, then poured onto the mould and cured at 80°C for 2 h. After curing, each PDMS slab was peeled off from the mould and cut to glass slide size (76 mm x 26 mm). The two slabs were superimposed and irreversibly bonded using oxygen-plasma bonding. Fluidic connections to the liquid flow channels and gas connections to the gas flow channels were made by punching holes through the PDMS using a tissue biopsy punch (Harris Uni-core, diameter 1 mm for liquid connections and 5 mm for gas connections).

**Figure 1.**
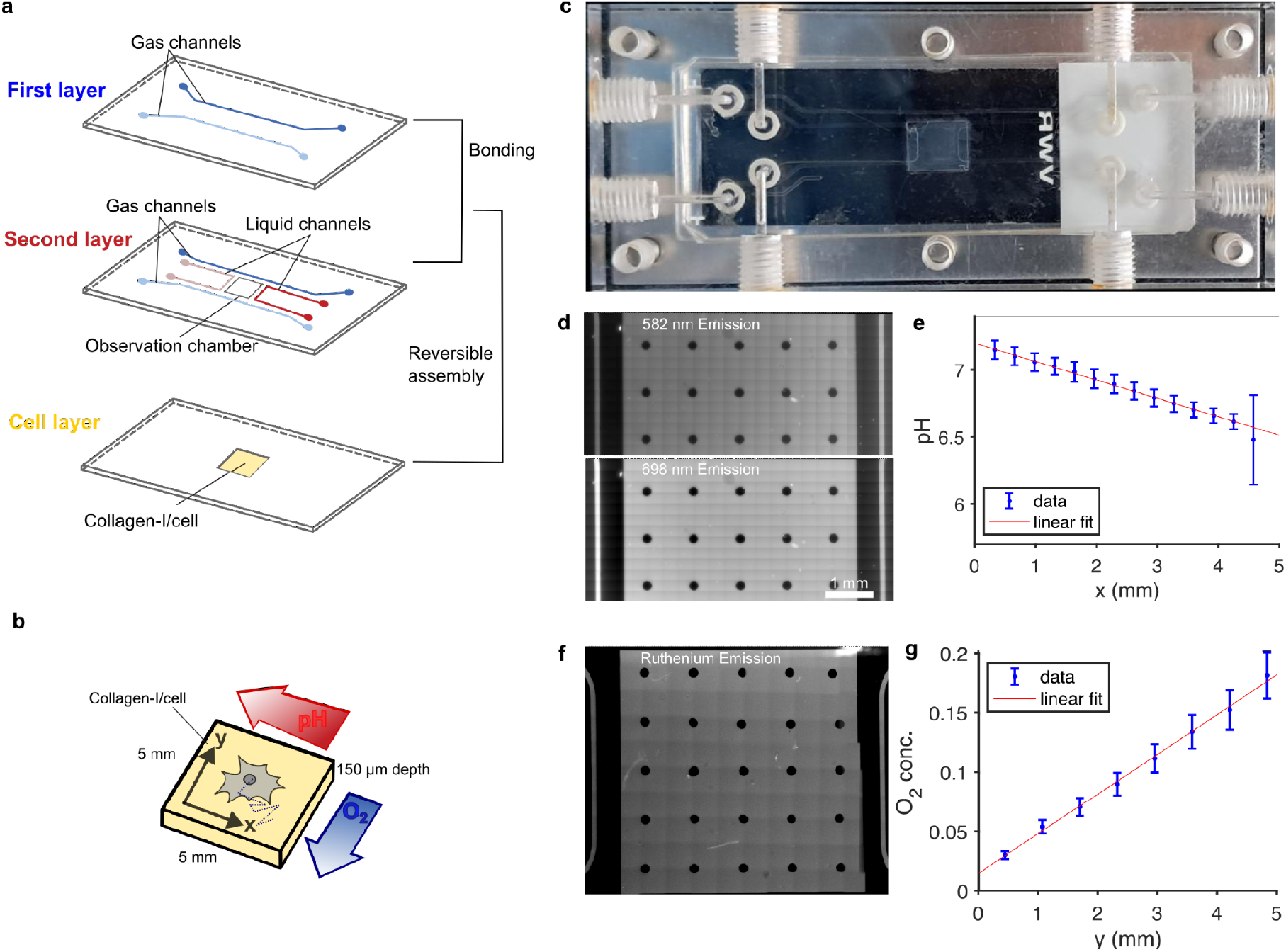
Microfluidic device and set-up. a) Schematics of the device assembly with two PDMS layers assembled through plasma bonding and a glass slide. b) The microfluidic device creates a pH gradient along the *x-*direction and an oxygen gradient along the *y*-direction over the collagen-I layer. c) Image of the device mounted on the microfluidic holder and connected to liquid and gas flow. d) Fluorescence signal of SNARF measured at steady state (4.5 h). e) pH values obtained from the ratiometric signal of SNARF and a calibration curve (see Figure S2). Error bars are obtained from the S.E.M of the measured intensity ratio and the slope of the calibration curve at the corresponding pH. f) Fluorescence signal of Ruthenium complex measured at steady state (14 h). g) Oxygen concentration obtained from the measured signal and the calibration at 1% and 20% oxygen. Error bars are calculated by error propagation of the signal standard deviation and the error on Kd.

In the top layer, the first pair of 500 μm-wide gas channels is defined (Fig. 1a). In the bottom layer, a 5 mm x 5 mm observation channel covers the gradient area, two liquid channels and two gas flow channels, symmetrically positioned to the gradient observation area (Fig. 1a). The liquid channels are 100 μm wide and 150 µm deep, separated from the gradient observation area by a 500 μm gap. The second pair of gas channels in the bottom layer are in plane with the gradient observation area. After assembling both PDMS layers, due to the thickness of the first PDMS layer, the pair of gas flow channels in the second layer is 0.8 mm away from the liquid flow channel and the edges of the gradient observation area. The pair of gas channels in the first layer is coplanar with the gradient observation area and placed 2 mm away from it to prevent gas channels from overlapping with the collagen layer, thereby increasing the robustness of the assembly (Fig. 1a).

The gas and liquid channels, as well as the gradient observation area, are separated by PDMS. Due to the high permeability of PDMS to oxygen, the oxygen tension in the gradient observation area can be controlled by adjacent gas- and liquid channels. Gas channels are supplied with either high (20%) or low (1%) oxygen (Fig. S1b). Away from the observation area, the gas and liquid channels run parallel to each other over a length allowing the solutions in the liquid channels (creating the pH gradient) to equilibrate at the high or low oxygen tension before they reach the region gradient observation channel.

### Lid with hydrogel cell culture unit

The second part of the device, which serves as a lid for the microfluidics channels, is a biological hydrogel unit for cell culture. It was prepared on a microscope glass slide with a frame of pressure-activated adhesive tape (adhesive, optical, MicroAmp) cut by CO_2_ laser-cutting (EpilogLaser). Its external dimensions of 76 mm x 26 mm fit a standard microscope slide and a window of 8 mm x 7.5 mm fits the position of the two liquid flow channels and gradient observation area on the PDMS device but do not overlap with the gas flow channels (red dashed box in Fig. 1b). The frame was laminated on a positively charged glass slide (Premium Printer Slides, VWR). A solution of 6 mg/mL collagen-I was prepared. For 200 µL of 6 mg/mL collagen-I solution, we mixed 20 µL of 10x PBS, 22.8 µL MilliQ water, 4 µL 1 M NaOH, and 153.2 µL of 10.45 mg/mL collagen-I (Rat tail Collagen-I, 734-1085, Corning). The collagen pH was controlled using pH paper (1.09543.001, Supelco) to achieve a pH of 7. The collagen-I solution was poured into the frame on a glass slide, covered with a thin Fluorinated Ethylene Propylene (FEP) foil, loaded with a glass slide on top to keep the FEP film flat, and allowed to gel at 37°C for 1 h. After gelation, the FEP film and the top glass slide were lifted, and the collagen-I film was kept in PBS, ready for use. The gel thickness, measured by imaging, is 150 µm.

### Device assembly and mounting

The two parts were held together with a homemade Poly(methyl methacrylate) (PMMA) holder, which comprised fluidic connectors and a metal frame. Screws to maintain the reversible sealing are adjusted at a maximum torque of 80 mN.m on 4 mm diameter screws. Pressure-based flow controllers (Fluigent, Flow EZ) were used to drive the liquid flow. First, the tubing connecting the flow controllers, flow rate sensors, and the device holder were pre-filled with solutions to remove all the air. Next, the oxygen plasma-treated PDMS device was placed on the top of the holder with all inlet- and outlet holes aligned. The PDMS microfluidic device was then covered with liquid before the glass slide with the collagen-I film facing down was assembled on top of the PDMS. The collagen-I film was aligned with the gradient observation area of the PDMS device and the gaps between gas flow channels and liquid flow channels (Fig. S1b, red dashed box). The pH of the solutions (5.7 and 7.6) was adjusted by adding the required amounts of HCl and NaOH to the cell culture media (DMEM, 41966-029, Gibco) and was subsequently equilibrated in the incubator (37°C, 5% CO_2_, 21% O_2_). Finally, the gas channels were connected. Low (1%) and high (20%) oxygen were injected into the gas flow channels by manually adjusting the overpressure, with an inlet pressure of 45 mbar corresponding to a flow rate of 13 µL/min.

### Numerical simulations

To adjust the dimensions of the microfluidic device and the experiment parameters, we simulated oxygen transport inside the PDMS and liquid using COMSOL Multiphysics 6.3. The gas and liquid flows in individual channels were simulated by solving the Navier-Stokes equations coupled with the mass continuity equation for an incompressible fluid. No-slip boundary conditions were applied to all channel walls. Both the transient and steady state of oxygen tension inside the device were computed by solving the convection-diffusion equation.

The simulation was achieved by combining the physics of *Laminar Flow* and *Transport of Diluted Species*. For the *Laminar Flow*, the inlets were defined as fully developed flow-based boundary conditions, along with the flow rates Q. Zero pressure and convection flux conditions were set at the outlets. For the *Transport of Diluted Species*, the inlets were defined as concentration constraint boundary conditions based on the oxygen concentration of supplied gases and solutions.

Based on Henry’s law, at the gas-PDMS boundaries within the PDMS device, the concentration of oxygen in the gas phase (*C*_*g*_) and PDMS (*C*_*p*_) is:

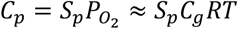

where *S*_*p*_ is the solubility of oxygen in PDMS, P_O2_ is the partial pressure of oxygen, *R* is the ideal gas constant, and *T* is the temperature. At the boundaries of PDMS and liquid, we have:

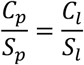

where *S*_*l*_ and *C*_*l*_ are the solubility and concentration of oxygen in liquid, respectively.

The boundary conditions between gas/PDMS and PDMS/liquid are described by the dimensionless partition coefficients *K*_*1*_ and *K*_*2*_, respectively (Fig. 2b).

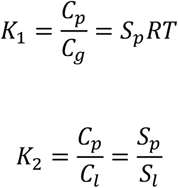

**Figure 2.**
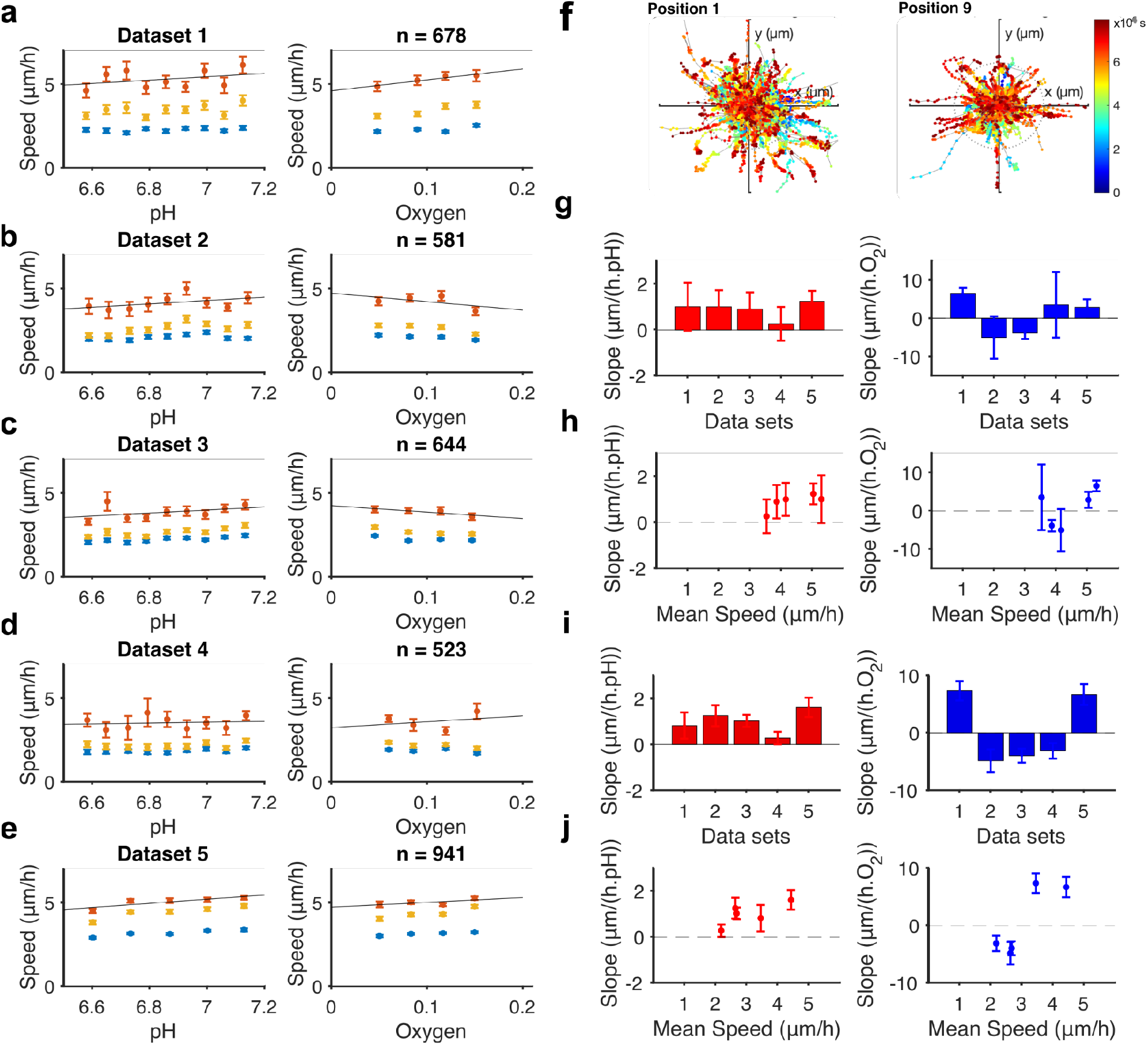
Cell motility results. a-e) Median speed as a function of pH and oxygen concentration for the ‘low’, ‘all’, and ‘high’ cells. Linear fit of the ‘high’ cells for five data sets. f) Example of cell tracks for 2 positions in dataset 5 corresponding to normal pH (position 1) and acidosis (position 9). g) Dependency of the cells’ median speed (in μm/h) with pH units and oxygen concentration displayed as the slope of the linear fit in (a-e). h) pH and oxygen dependency on the average cell motility (error bars are the standard error of the slope). i-j) same as g-h) for all cells.

In the simulation, the liquid is assumed to be equilibrated with air until it reaches the inlet of the device, that is, the oxygen concentration is 21%. We also assume that the initial oxygen tensions in the liquid of the gradient observation area and PDMS are 21% at the onset of the simulation. The oxygen tension at the top and bottom faces of the PDMS was not considered since the PDMS slab is held between two gas-impermeable surfaces (PMMA and glass). Water is used as the liquid in the liquid channels. The physical properties of each material (water, PDMS, and gas) were selected directly from the materials library. The solubility of oxygen in PDMS (*S*_*p*_) and water (*S*_*l*_) are 1.25 mM/atm and 0.218 mM/atm ^13^, respectively. Input data for the simulation are listed in Table S1.

### pH gradient and calibration curve

The pH gradient in the device was visualized and quantified using 40 µM of 5-(and-6)-Carboxy SNARF-1 (#C1270, ThermoFisher). Ratiometric measurement was performed using excitation at 488 nm and emission filters of 582/75 nm and 698/70 nm. The ratio was measured along the *x*-direction of the observation area.

To determine the pH values across the observation area, a calibration curve was generated in a separate experiment. The SNARF ratiometric signal was determined as a function of pH, using six Ringer solutions (115 mM NaCl, 5 mM KCl, 1 mM Na_2_HPO_4_, 1 mM CaCl_2_, 0.5 mM MgCl_2_ and 24 mM NaHCO_3_) of known pH. In this calibration experiment, separate collagen-I films were soaked overnight at their respective pH before being mounted in the device and measured under the appropriate pH solutions and oxygen flow. The calibration and gradient experiments in the absence of cells were performed at 20°C.

### Oxygen gradient and calibration curve

The oxygen gradient in the microfluidic system was measured using the oxygen-sensitive dye tris(2,20-bipyridyl) dichlororuthenium (II) hexahydrate (RTDP) (Sigma-Aldrich #544981). The fluorescence intensity of the dye is quenched in the presence of oxygen ^26^.

RTDP was dissolved in PBS at a concentration of 5 mg/mL. Collagen-I films prepared on glass slides were soaked in RTDP solution overnight to equilibrate the RTDP completely and ensure that fluorescence intensities solely represent the oxygen gradient. The assembly process was as described above (‘*Device assembly and mounting’*), except that all PBS solutions, including those supplied to the liquid channels, contained RTDP. At the onset of the experiment, both liquid channels were connected and a flow rate of 3 μL/min was applied. Both gas channels were connected to 20% oxygen for 30 min and a reference image of the RTDP signal at 20% oxygen was then recorded. Next, the gradient was established by connecting the gas channels to 20% oxygen and 1% oxygen. The RTDP fluorescence intensity in the observation area was recorded using time-lapse imaging at 4, 8, 12, and 14 h, with a filter cube consisting of an excitation filter at 480/30 nm, a dichroic mirror at 565 nm, and an emission filter at 630/69 nm (AHF analysentechnik). Finally, all gas channels were connected to 1% oxygen to record the second reference image. Using the pair of reference images at 1% oxygen and 20% oxygen, we measured a K_d_ of 1.346 ± 0.003 so that the RTDP intensity in the observation area can be converted to oxygen concentration using:

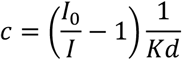

with *I*_*0*_ is the intensity in the absence of quenching at 0% oxygen, evaluated from the calibrated value of Kd, *I* is the measured intensity, and *c* is the oxygen concentration. Both the calibration and the experiment were performed at 20°C.

### Monitoring cell motility

#### Protocol for cell culture outside the chip

MDA-MB-231 cells were cultured in DMEM (41966-029, Gibco) in an incubator with 5% CO_2_ and 21% O_2_ and split twice a week. Cells were maintained in culture until passage 20.

#### Protocol for the cell culture in the chip

MDA-MB-231 cells were detached using trypsin (T4174, Sigma) and counted to achieve a final concentration of 4 × 10^6^ cells/mL in collagen at 6 mg/mL. Cell viability was checked during cell counting and was about 95% for each experiment. After counting, cells were centrifuged at 300 g for 3 min to form a cell pellet and then resuspended in freshly prepared collagen (6 mg/mL). Collagen with embedded cells was stored in DMEM media in the incubator for 72 h before the microfluidic experiment.

Imaging: We imaged the cells in a z-stack consisting of 25 planes at a 5 μm pitch, using bright field at 20x magnification (Nikon, 20x/0.70) on an inverted microscope (Nikon Eclipse Ti2-E) equipped with a motorized stage and a temperature/CO_2_-controlled environmental chamber (OKOlab, UNO-T-H-PREMIXED). The temperature of the collagen-I unit was measured in a separate experiment, exploiting the temperature sensitivity of RTDP ^26^. This showed that the temperature in the collagen is within a few degrees of 37 °C (Fig. S8). We imaged cells on a 9 × 4 grid of 36 fields of view in the gradient observation area (Fig. S6). The camera (DS-FI3, Nikon) has a pixel size of 4.8 μm. Each field of view is 1440 × 1024 pixels (345 × 245 μm). For each position, we recorded a time-lapse (30 min) for 22 h. Data set 5 was recorded using a different camera (Teledyne, Kinetix), with a field of view of 1200 × 1200 pixels (390 × 390 μm). We imaged cells at 20 fields of view, arranged on a 5 × 4 grid of positions in the gradient. For each position, we recorded a time-lapse video, acquiring images every 10 min for 22 h.

Cell motility was tracked on a 2D projection of the z-stack using Trackmate and Cellpose software as previously described ^12^. Briefly, a Cellpose ^27^ segmentation model was trained on a 2D projection of the z-stack. For the data sets with the 1440 × 1024 pixel field of view (data sets 1-4), the image was resampled to a 1077 × 766 pixels field of view to obtain a pixel size of 0.32 μm, equivalent to that of data set 5. In Trackmate ^27,28^, we generated tracks that allowed no track splitting and no gaps. Using a separate, simpler tracking on a single *z*-plane, we measured drift in each field of view, taking into account stage *xy* drift and gel deformation. This local drift was then subtracted from all tracks measured in this field of view. We selected tracks based on the maximum distance traveled by the cells from their first position, accepting a 10 μm distance as the minimum for a viable cell. Of those tracks, we also categorized high and low motility cells based on their mean speed, using a threshold of 5 μm/h.

### Statistical analysis

In each field of view, the median speed of each viable cell track is calculated. From this, we calculate the mean and standard error of the mean of the cells’ median speed per field-of-view and assign a value of pH and oxygen corresponding to the center of the field-of-view in the gradients (linear fits in Fig. 1e and g). We calculate the slope of the speed versus pH curve and speed versus oxygen curve using linear fits of the data and report the standard error of the slope as error bars. All analysis is conducted in MATLAB (R2024a, MathWorks).

## Results

### Overall device design and operation

We developed a microfluidic device that creates orthogonal pH and oxygen gradients (Fig. 1). The device consists of two parts: a PDMS microfluidic chip and a removable lid featuring a collagen-I hydrogel layer suitable for cell culture. The PDMS chip itself is built of a bottom layer that comprises a 5 × 5 mm observation area with a microfluidic channel on two opposite sides. The 100 × 150 μm microchannels are separated from the observation area by a 500 μm PDMS gap and convey culture media at different pH. In the same layer, a pair of gas channels is added, separated from the edges of the observation area by 500 μm. A second PDMS layer defines another pair of gas channels running on top of the first one. Both layers are fabricated via PDMS casting and bonded via oxygen plasma bonding. This PDMS device is pressed against a glass slide comprising a collagen-I layer so that the observation area and a portion of the microchannels cover the collagen-I layer on the glass slide. Meanwhile, the connection holes of the PDMS device align with the liquid and gas connections of the holder.

Upon assembly, the microchannels and the observation area are connected through two hydrogel barriers, allowing diffusion but preventing the convection flow of the liquid. Using two solutions at different pH in the microchannels, a pH gradient is established by diffusion through the hydrogel barriers. Simultaneously, gas channels are supplied with either high or low oxygen gas. Due to the gas permeability of the PDMS, a gradient of oxygen is established across the whole device through the PDMS, the liquid, and the hydrogel. At steady state, the concentration gradients of hydronium ions and oxygen are expected to be linear in the observation area along the *x*- and *y-directions*, respectively, i.e., orthogonal to each other.

### Calibration experiments validate the orthogonal pH and oxygen gradients

We first validated the linearity of the concentration gradients in the observation area using fluorescent pH- and oxygen indicators in solution. The gradients were generated with Ringer solutions of pH 5.7 and 7.6 and oxygen concentrations of 20% and 1%, respectively, regardless of whether the pH- or the oxygen gradient was studied.

To assess the pH gradient linearity, we measured the fluorescence intensity of the pH-sensitive dye SNARF-1 (Fig. 1c). The data show that a pH gradient is established in the x-direction after 1 h and remains entirely linear after 4.5 h. This is as expected from the estimate of the time to gradient using the total 6 mm distance between the liquid channels (1 h estimated from L^2^/D, where L is the distance between the source and sink, and D is the diffusion coefficient of hydronium ions in water). Fluorescence intensities were converted to pH using a calibration curve generated in a separate experiment (Fig. S2a), showing that the pH gradient is linear. Hydronium ion concentrations were calculated from the pH values in the gradient, showing that the concentration gradient is also linear within the confident calibration interval of SNARF-1. A linear fit to the pH gradient indicates that the pH range accessible in the observation area is 6.5 - 7.2, with a slope of 0.137 ± 0.002 units per mm. Conversely, the ratiometric signal, pH, and hydronium ion concentration all vary linearly with the position along the *x*-axis within the error bars of our measurements. In a separate experiment, we monitored the 582 nm SNARF-1 value for up to 22 h and confirmed that the gradient remains stable once established (Fig. S2). Both experiments were performed at room temperature (20°C).

For probing the oxygen concentrations, we used the fluorescent Ruthenium complex RTDP ^26^ (Fig. 1e-f and Fig. S4). As expected, an oxygen gradient was established in the *y*-direction in the observation area, while there was no gradient in the *x*-direction (i.e., along the pH gradient) (Fig. S4d). The oxygen gradient remained stable after 1 h and persisted for up to 18 h, whereas numerical simulations and other control experiments suggest that the gradient should establish within 90 min (see Fig. S4) of the onset of gas flow in the sink and source channels. Here, we measure the usable range of oxygen tension to be from 1.5 to 18.2% oxygen when the source and sink concentrations are 1% and 20%, respectively, using an independent calibration of the fluorescence indicator’s response to oxygen at a constant temperature. The slope of the gradient is 3.3 ± 0.7% per mm.

The measured usable range is in very good agreement with numerical simulations, which predict a usable oxygen range in the collagen-I layer of 2.5% to 18.1% over the 5 mm width of the observation area.

Taken together, these results show that our microfluidic device can generate stable orthogonal pH and oxygen gradients within a few hours after mounting. Within the observation chamber, the gradients are linear, ranging from pH 6.5 to 7.2 and from 1.5% to 18.2% oxygen, respectively.

### The pH and oxygen gradients differentially impact cancer cell motility

Cancer metastasis relies on the acquired ability of cancer cells to migrate away from the primary tumor and invade through the tumor-intrinsic and -surrounding extracellular matrix, intravasate into the vasculature, and eventually extravasate to colonize new niches ^29^. The migration and invasion of cancer cells are highly dependent on environmental pH ^1,2,30^, as well as on oxygen concentrations ^31^. However, the combined effect of specific pH and oxygen values has, to our knowledge, never been studied.

To investigate how orthogonal gradients of pH and oxygen influence cell motility, we seeded MDA-MB-231 triple-negative human breast cancer cells in collagen-I on glass slides and allowed them to recover for 72 h at 21% O_2_/5% CO_2_ in a 37°C incubator. On the morning of the experiment, the device was assembled and mounted with liquid and gas tubing in the device holder on the microscope stage. Images were acquired periodically over multiple fields of view for 22 h (see section ‘Monitorin*g cell motility’*). After CellPose-aided cell detection, we quantified motility at a given pH- and oxygen value by determining the median speed of each cell and pooling data for all cells imaged at the same pH position (Fig. 2a-e). Consistent with our previous study using a device producing a pH gradient alone ^12^, the cell median speed was highest at normal tissue pH (7.4), decreasing with decreasing pH. In the orthogonal direction, along the oxygen gradient, motility tended to be highest at the highest oxygen concentration (Fig. 2f-g). However, in experiments where the average motility was low (3 data sets, Fig. 2f-g), this pattern was not observed (Fig. S7f-g). These results show that human cancer cells move in a pH- and oxygen-dependent manner in a collagen-I matrix.

## Discussion

Complex microenvironmental gradients play key roles in a wide range of physiological and pathophysiological processes, from embryonic development over kidney function to cancer ^4,32,33^. Here, we present the first microfluidic device capable of generating precise orthogonal gradients of pH and oxygen concentrations while enabling live imaging of cells embedded in a biological hydrogel. Notably, we achieve this while retaining the reversible sealing capability that proved crucial to the integration with a spatial transcriptomics workflow, as demonstrated in a previous work ^12^. Using similar approaches, the workflow can also be integrated with other spatial ‘omics techniques.

We successfully determined the pH gradient experimentally, discovering that the concentration gradient between the source/sink and the edges of the observation area was steeper than within it (Fig. S5 a and c). This is expected since in the gap between the liquid channels and the observation area, 50% of the height is occupied by the PDMS barriers, resulting in a reduced flux of ions.

However, we also noted two quantitative discrepancies with expectations only considering a one-dimensional concentration gradient of hydronium ions 1) the time scale of the establishment of a linear gradient in the dual gradient device was significantly longer than what is expected and 2) the pH drop in the gap between the observation area and the source/sink channels are asymmetric. The main reason for the first discrepancy may be the diffusion coefficient of hydronium ions in collagen-I. The diffusion coefficient of salts can be assumed to be the same in hydrogels and water; however, a reduced diffusion coefficient of hydronium ions in hydrogels has been suggested previously ^34^. Still a stable gradient is established quickly compared to the time needed to study cell motility using a device architecture that makes the gradient less dependent on flow rates than the tree-like mixer devices ^12,21^.

Regarding the second discrepancy, because hydrogel barriers of equal width will cause equal concentration differences, a reduced accessible range of pH compared to the source and sink pH values is expected. However, the measured pH drop was highest at alkaline pH values. Both observations suggest that predicting the correct pH gradient requires the inclusion of the equilibrium reactions of buffering by HCO_3_^-^/CO_2_ and the relevant organic buffer systems in the growth medium in the model. This has to our knowledge so far not been done, and our current efforts are aimed at integrating this in future models.

To aid in the development of the design and to gain a deeper understanding of the dynamics of the gradients, we simulated the oxygen conditions using COMSOL modeling and compared the results to quantitative experimental analyses. It is noticeable that, in the case of the oxygen gradient, the slope of the gradient outside the observation area is smaller than that inside, due to the higher diffusion coefficient of oxygen in PDMS compared to water/hydrogel (Table S1). As a result, the part of the gradient located in the gaps between the observation area and the gas channels is minimal, and the accessible oxygen tension range is maximized (Fig. S5 b and d). In our experiment with cells, we supply 20% and 1% oxygen in the source and sink. We measure the oxygen concentration range under those conditions using our calibration values measured at 1% and 21% and obtain a range of 1.5-18.2% (Fig. S5d). Numerical simulations predict a range of 2.5% to 18.1%. Here, we note that the width of the gap (500 μm) between the gas channels and the edge of the observation area accounts for the majority of the remaining few percent at both the low and high ends of the range. The oxygen range in the observation area, from 1.5% to 18.2%, comprises hypoxic conditions for cultured cells that are normally maintained at 21% oxygen. However, because physiological oxygen conditions in our tissues are generally substantially lower than 21% and can be essentially 0% in certain conditions, such as poorly perfused tumor regions ^35^, it would be desirable to further decrease oxygen tension.

Achieving concentrations of oxygen below 1% is a general and important challenge in microfluidic systems ^13,15,25^, given that oxygen concentrations especially in pathophysiological conditions can reach levels very close to anoxia. In conditions where 0% and 21% are used as sink and source (for example, using nitrogen and air), this would produce an accessible range only slightly wider (Fig. S5b). A narrower but lower oxygen tension range can be attained by decreasing the oxygen concentration at the source without changing the device design. The gradient would only be weaker. Numerical simulations suggest that using 10% oxygen at the source produces a 0.7% - 9.7% range. A limit to attaining a lower oxygen tension is the rediffusion of the oxygen from the surroundings into the PDMS through the edge of the device. Our numerical simulations show that preventing the edge of the PDMS device from being in contact with air would reduce the lower limit of the usable oxygen range to 0.5%. This could be done by infusing nitrogen into the microscope stage.

We quantify the motility of the cells as the median speed and assign a pH and oxygen value in the gradients (Fig. S6). Others ^36^ have measured an increase in motility as the increased number of cells populating an artificial ‘wound’ and found that motility increased with pH, at constant pH gradient values higher than ours (lowest is 0.2 units/mm). By comparison, in our assay, the pH and oxygen gradients are superimposed but not collinear, and consequently, the effect of each gradient can be disentangled. Indeed, to our knowledge, ours is the first study to address cancer cell migration in such superimposed gradients, which more closely resembles biological conditions, such as those found in tumors.

While our device has the capacity to support breakthroughs in understanding how complex TME gradients impact cell behavior, gene expression and metabolism, the current design can be further improved in future iterations. A key perspective is to make the design more scalable and convenient, enabling off-the-shelf use. A current limitation is that the collagen-I hydrogel needs to be relatively stiff (6 mg/mL) for the device to perform reliably. While this corresponds well to conditions in tumors ^37^, it would be advantageous to develop materials that are compatible with biological hydrogels with a wider range of stiffnesses, given the importance of mechanical forces for cellular phenotypes. There is also potential to further improve the cell tracking workflow with more advanced 3D segmentation AI tools as they become available.

## Conclusion

We designed and validated a microfluidic device that enables the generation of stable, orthogonal pH and oxygen gradients across an observation area in which cells are cultured in a biological hydrogel. We characterize the system by combining numerical simulations with experimental validations of the behavior of the gradients. While we demonstrate the use of the device for analyzing how pH and oxygen gradients impact cell motility, the system is compatible with a wide range of other phenotypic analyses and can be adapted for generating other microenvironment gradients.

## Data availability

All data supporting the findings of this study are presented within the manuscript and its online supplement. Any additional information can be obtained from the authors upon reasonable request.

## Author contributions

Conceptualization, AS, SFP, RM; methodology, YL, LDS, MRC, RM; formal analysis, YL, LDS, MRC, RM; investigation, YL, LDS, MRC; data curation, YL, LDS, RM; writing—original draft preparation, RM, SFP; writing—review and editing, YL, LDS, MRC, JA, AS, RM, SFP; visualization, YL, LDS, MRC, RM; supervision, RM, AS, SFP; project administration, RM, SFP; funding acquisition, RM, AS, SFP. All authors have read and agreed to the published version of the manuscript.

## Acknowledgments

The project was funded by the Novo Nordisk Foundation (NNF19OC0057739 and NNF21OC0069598 to SFP, RM, and AS), the Carlsberg Foundation (CF20-0491 to SFP).

