## Supplemental Material for "A microfluidic model to recapitulate pH and oxygen gradients in solid tumors"

**List of supplementary materials**

Table S1. Parameters used for numerical simulations.

Figure S1. Device design and experimental set-up.

Figure S2. Calibration of pH using SNARF fluorescence.

Figure S3. Numerical simulation model and results of oxygen gradients in the device.

Figure S4. Calibration of oxygen using RTDP fluorescence.

Figure S5. pH and oxygen gradient slopes outside the observation area.

Figure S6. From time-lapse imaging data to cell tracking data.

Figure S7. Cell tracking results for high and low motility cell populations.

Figure S8. Chip temperature measurement using RTDP.

**Table S1.** Parameters used for numerical simulations.

| | | | |
|---|---|---|---|
| Wm | 5 [mm] | 0.005 m | Gradient area width |
| Hm | 5 [mm] | 0.005 m | Gradient area length |
| Wl | 40 [mm] | 0.04 m | Liquid flow channel length |
| Hl | 100 [μm] | 1E-4 m | Liquid flow channel width |
| Wg | 40 [mm] | 0.04 m | Gas flow channel length |
| Hg | 500 [μm] | 5E-4 m | Gas flow channel width |
| Dp | 4e-9 [m²/s] | 4E-9 m²/s | Diffusion coefficient, $O_2$ in PDMS |
| Dl | 2e-9 [m²/s] | 2E-9 m²/s | Diffusion coefficient, $O_2$ in water |
| Dg | 2e-5 [m²/s] | 2E-5 m²/s | Diffusion coefficient, $O_2$ in gas |
| Hc | 150 [μm] | 1.5E-4 m | Channels height |
| Qg | 18000 [μL/min] | 3E-7 m³/s | Gas flow rate |
| Ql | 2 [μL/min] | 3.3E-11 m³/s | Liquid flow rate |
| T | 37°C | 310.15 K | Temperature |
| Sp | 1.25 mM/atm | 1.23E-5 s².mol/(kg.m²) | Solubility of $O_2$ in PDMS |
| Sl | 0.218 mM/atm | 2.15E-6 s².mol/(kg.m²) | Solubility of $O_2$ in water |

* Temperature-dependent values are reported at 37°C [1].

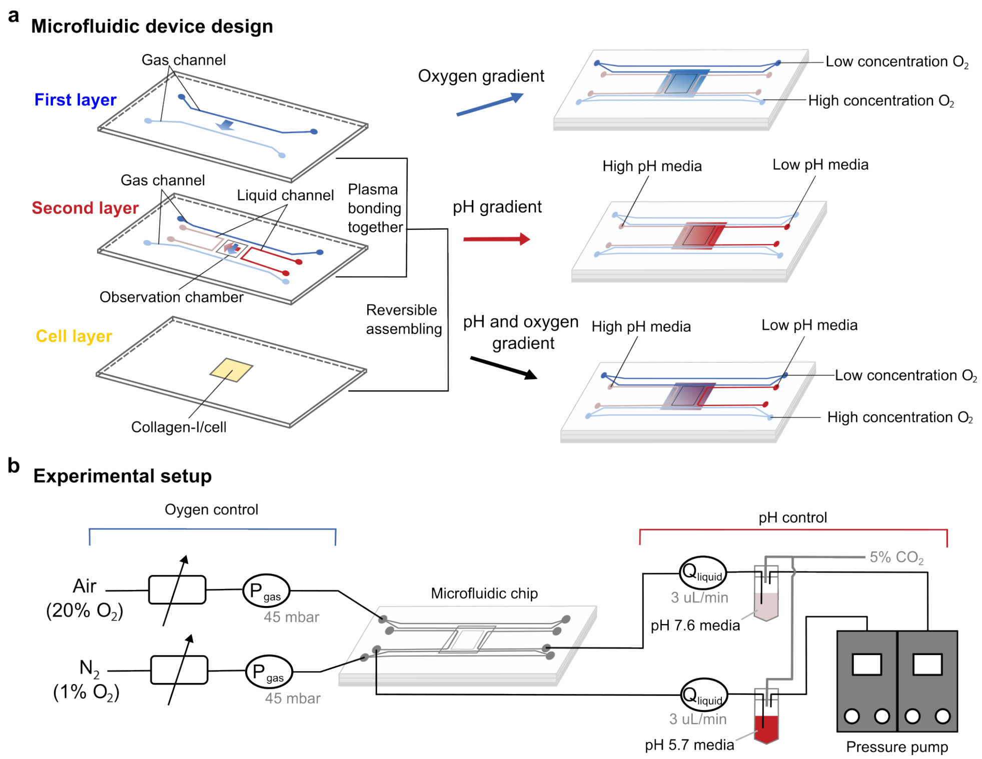


Figure S1. Device and set-up. a) The device is made of three layers. The first two layers, made of PDMS and assembled irreversibly through plasma bonding, comprise two pairs of gas channels and one pair of liquid channels. The PDMS assembly is reversibly sealed with a glass slide comprising a collagen-I layer with cells. b) Oxygen is supplied to the gas channels at 20% and 1%, creating an oxygen concentration gradient over the collagen-I film. pH solutions at 5.7 and 7.6 are supplied to the liquid channels, thus creating a pH gradient over the collagen-I film. Due to the device geometry, the oxygen and pH gradients are orthogonal to each other.

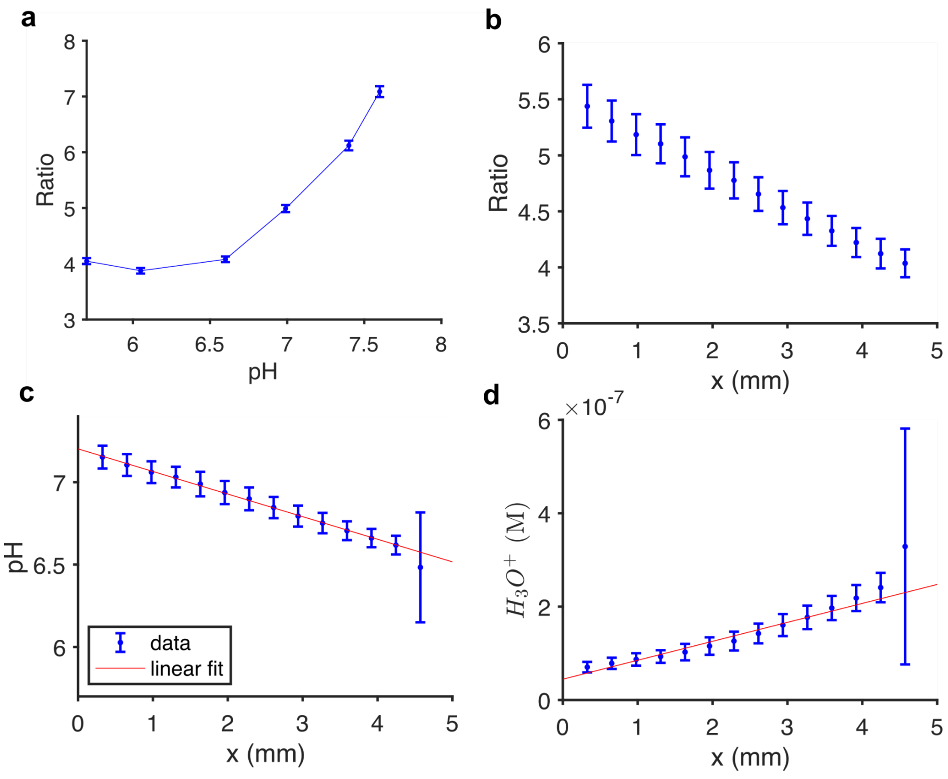


Figure S2. Results of pH gradient characterization using the ratiometric fluorescence signal of SNARF-1. a) A calibration curve of the ratiometric signal using the emission at 582 nm and 698 nm. b) Ratiometric signal measured in the observation area of the device along the *x*-direction (mean and standard deviation). c) pH values obtained by linear interpolation on the calibration curve. The fit is used to calculate the accessible pH range of pH 6.5 to pH 7.2 with a slope of 0.137±0.002 pH units per mm. Error bars are propagated from the error on the ratiometric signal. d) Hydronium ions concentration as a function of the distance along the *x*-direction calculated from the pH values (fit in red). Error bars are propagated from the error on the pH.

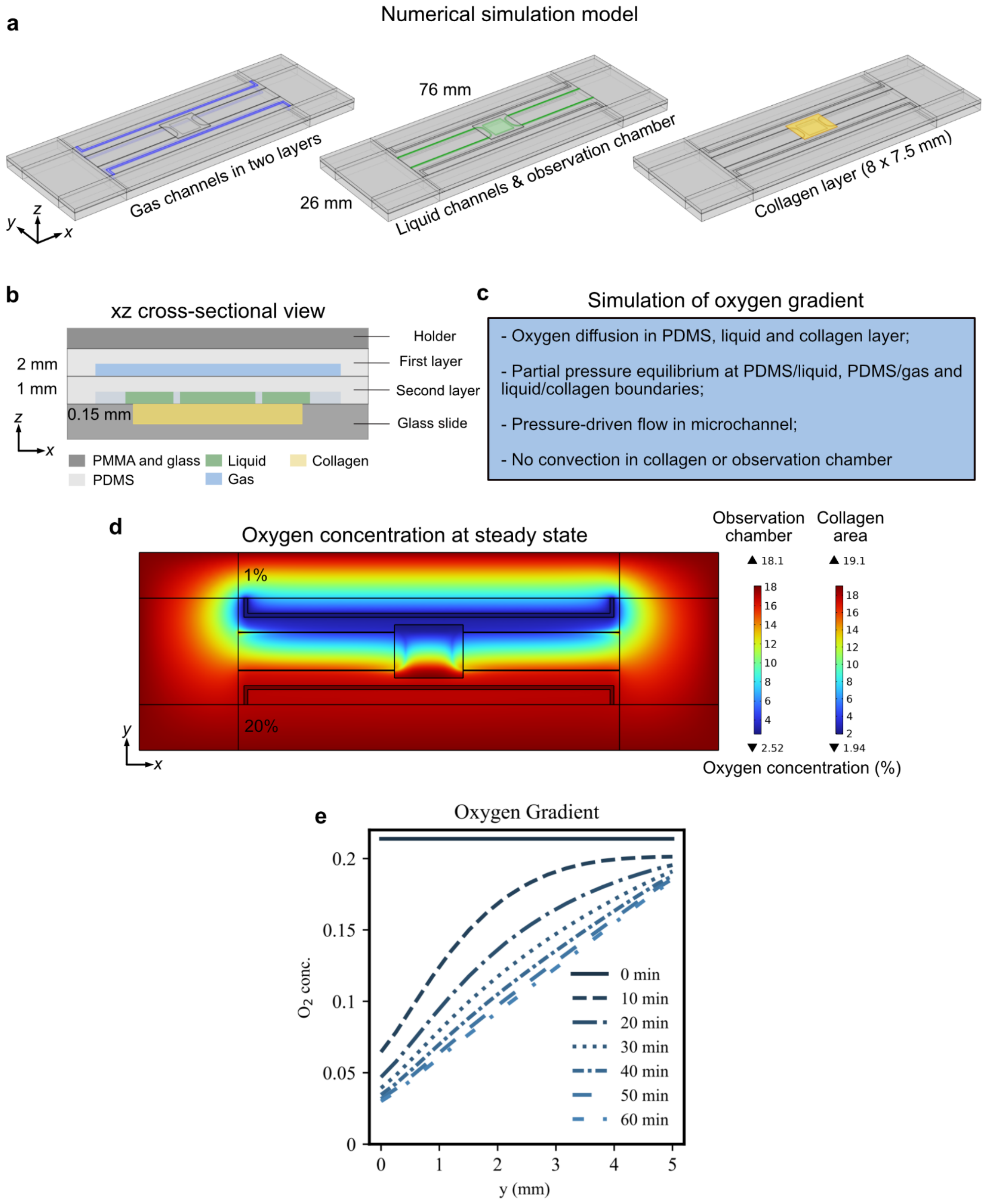


Figure S3. Results of numerical simulations of the oxygen gradient. a) Geometry of the numerical simulation model. b) Sketch of the model cross section showing the overlap of the liquid channels (100 µm depth) and observation area with the collagen layer (150 µm depth, not to scale). c) Brief implementation notes for Comsol's Transport of Diluted Species (tds) module. Numerical values for the simulation parameters are displayed in Table S1. d) *X-y* view of the oxygen concentration at steady state in the chip, including the concentration range in the observation chamber and collagen layer. e) Oxygen concentration in the collagen layer along y and within the observation area (5 mm) as a function of time. The time interval for oxygen profile simulations is 10 min.

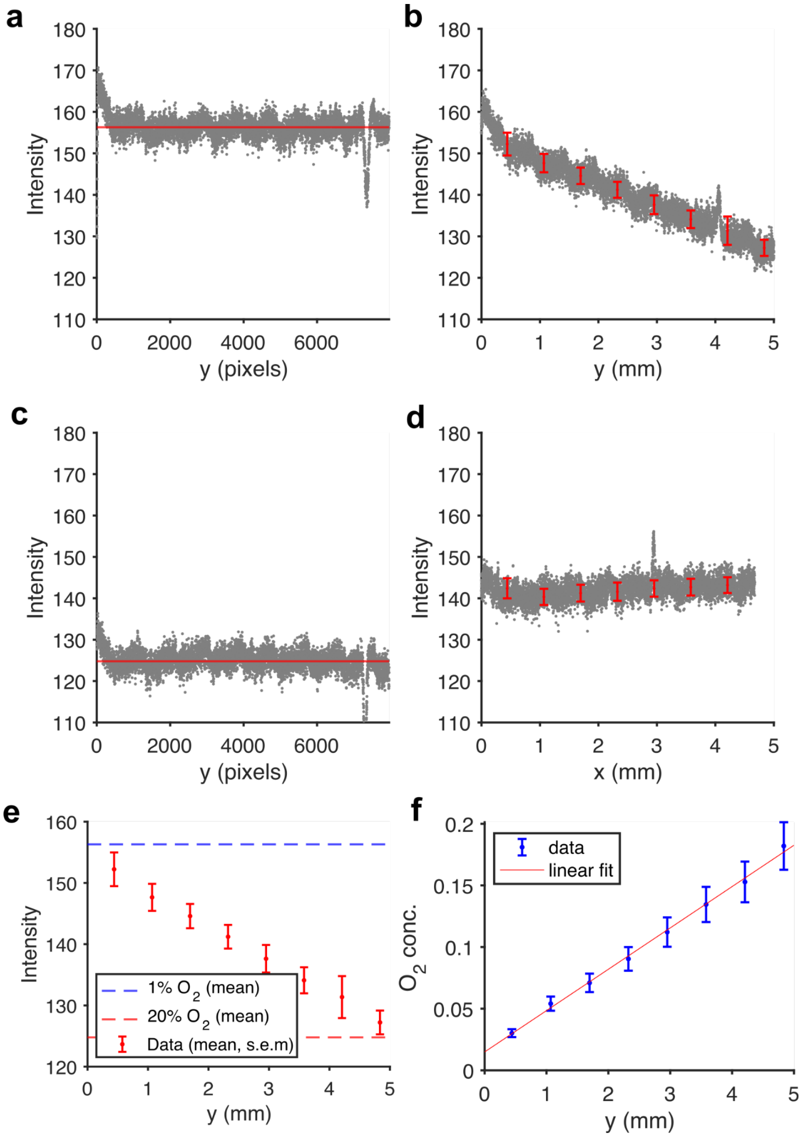


Figure S4. Oxygen gradient characterization using Ruthenium complex. a) Fluorescence intensity along the *y*-direction in the observation area for the low oxygen calibration (all gas channels at 1% oxygen). b) Same for the high oxygen calibration (all channels at 20% oxygen). c) Same after 14 h gradient conditions. Blocked intensity values are shown as mean and s.e.m. d) Fluorescence intensity along the *x*-direction showing a constant signal (no gradient). e) Plot showing the calibration values (124 and 156 units, respectively) and the intensity values along the gradient in the *y*-direction. f) Oxygen concentrations calculated from the blocked intensity values and the calibration values. Error bars are propagated from the s.e.m of each block. The linear fit indicates a slope of 3.3±0.7% oxygen concentration per mm.

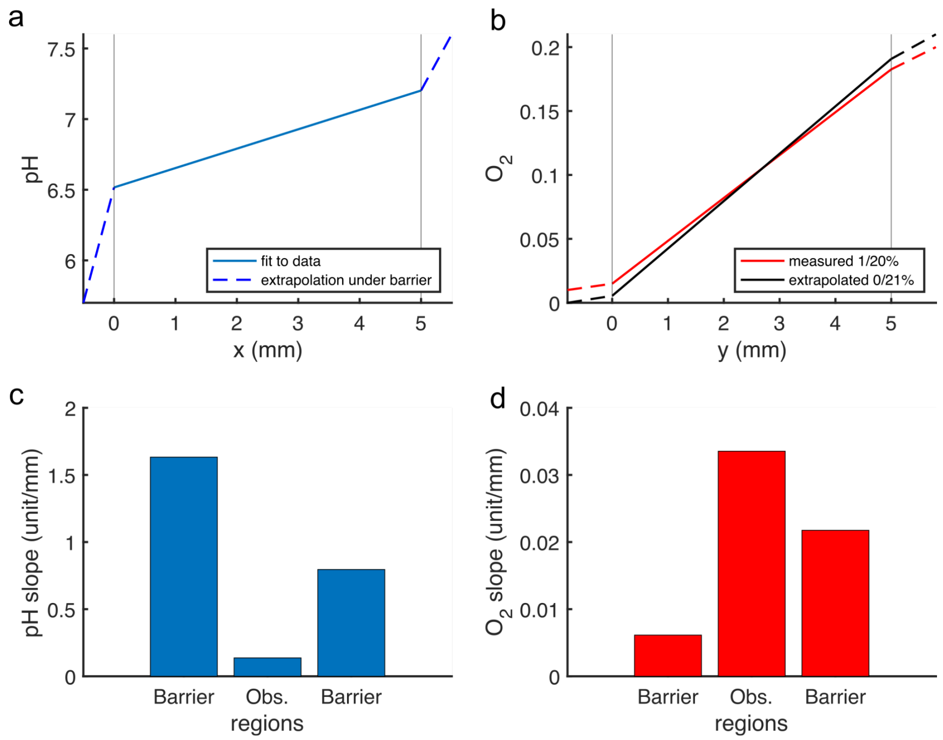


Figure S5. Extrapolation of the pH and oxygen gradient values outside the observation area. a) The pH gradient in the hydrogel barriers separating the gradient observation area from the source (pH 7.6) and the sink (pH 5.7) is steeper than in the observation area. b) The oxygen gradient extrapolated outside the observation area is less steep than in the observation area (red line). The oxygen gradient is extrapolated for conditions where the sink/source is 0%/21% (black line), assuming a constant ratio between the slopes in the different regions. c) pH gradient slopes in the different regions showing reduced ion flux in the barrier regions. d) Oxygen gradient slopes in the three regions. The data is consistent with a higher diffusion coefficient in the PDMS compared to water.

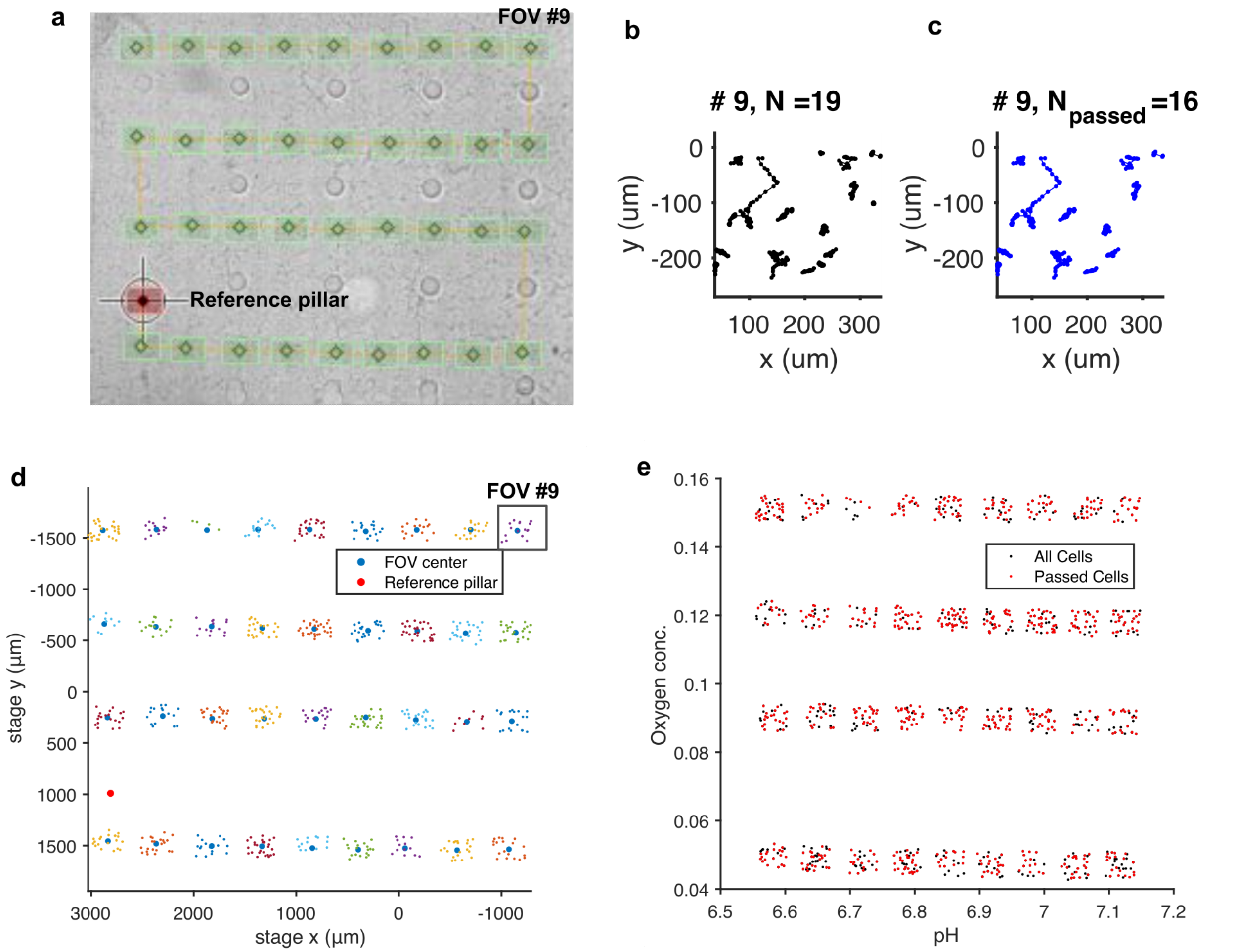


Figure S6. Field of views, cell tracks, and sampling of the gradient. The example is from data set 1. a) Overview of the field of view positions (green) overlay with a low-resolution image of the collagen-I gel. The position of the FOV within the observation area of the device is referenced using an image of a pillar (red). b) Example of cell tracks of N>20 positions for the field of view #9 after drift correction. c) Cell tracks of N>20 and a maximum distance traveled larger than 10 µm for the same field of view. d) Mean positions of the cell tracks shown in absolute stage positions and colored by field of view. The cell's global position is calculated from the position in the field of view and the stage position of the center of the corresponding field of view (blue dots). e) pH and oxygen values are assigned to each field of view based on the center of the field of view. Here, the positions of the cells in the double gradient are also shown.

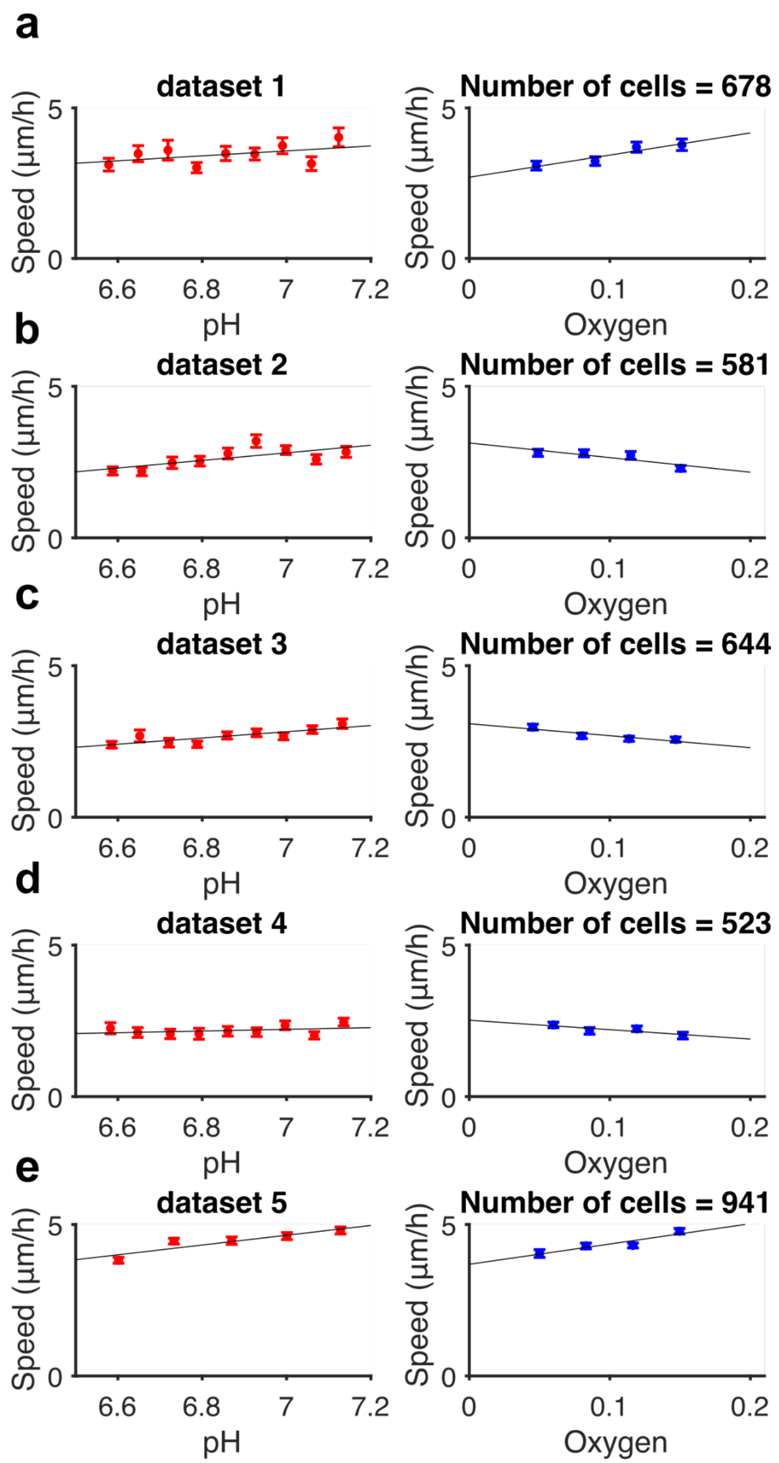


Figure S7. Cell tracking: analysis of cell median speed for all cells in data sets 1-5. Linear fits are used to measure the cells' median speed dependence on pH and oxygen, as shown in Figures 2h-i of the main text.

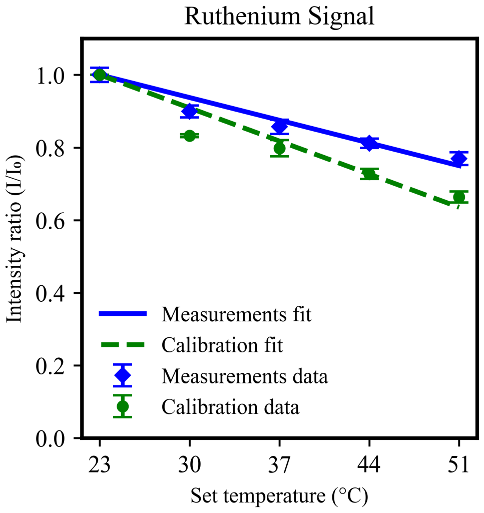


Figure S8. Calibration and measurement of the temperature using the RTDP fluorescence signal. The calibration data (mean and s.e.m.) were acquired using a 5 mM solution mounted on a microscope glass slide and a cover slip, ensuring optimal temperature control using the stage enclosure (see 'Materials and Methods'). The fluorescence signal is normalized to the value at room temperature (23°C). At 37°C, the fitted intensity ratio is 0.875. The temperature in the observation area of the device is estimated to be 32.6°C based on a linear fit to the measured intensity ratio (mean and s.e.m).
